# The Functional Connectivity between the Locust Leg Pattern Generating Networks and the Subesophageal Ganglion Higher Motor Center

**DOI:** 10.1101/226167

**Authors:** Daniel Knebel, Jan Rillich, Leonard Nadler, Hans-Joachim Pflueger, Amir Ayali

**Author notes:** These should be considered joint first author (Daniel Knebel, Jan Rillich). Author for Correspondence: Amir Ayali.

## Abstract

Interactions among different neuronal circuits are essential for adaptable coordinated behavior. Specifically, higher motor centers and central pattern generators (CPGs) induce rhythmic leg movements that act in concert in the control of locomotion. Here we explored the relations between the subesophageal ganglion (SEG) and thoracic leg CPGs in the desert locust. Backfill staining revealed about 300 SEG descending interneurons (DINs) and some overlap with the arborization of DINs and leg motor neurons. In accordance, in *in-vitro* preparations, electrical stimulation applied to the SEG excited these neurons, and in some cases also induced CPGs activity. Additionally, we found that the SEG regulates the coupling pattern among the CPGs: when the CPGs were activated pharmacologically, inputs from the SEG were able to synchronize contralateral CPGs. This motor output was correlated to the firing of SEG descending and local interneurons. Altogether, these findings point to a role of the SEG in both activating leg CPGs and in coordinating their oscillations, and suggest parallels between the SEG and the brainstem of vertebrates.

## Introduction

A longstanding and fundamental question in neuroscience is that of how the brain coordinates motor behaviors (Penfield and Boldrey, 1937; Schmidt and Lee, 2005; Fritsch and Hitzig, 2009). Ample research across species and methodologies has revealed a differentiation between three levels of motor control: central pattern generators (CPGs - neuronal oscillators that can produce rhythmic motor output in the absence of sensory or descending inputs; see recent reviews in Marder and Bucher, 2007; Mulloney and Smarandache, 2010; Marder, 2012; Rybak et al., 2015); sensory feedback; and higher motor centers. In most cases, the CPGs determine rhythmic alternating activity of specific antagonistic muscles. The CPGs by themselves, or often together with a few of the many sensory feedback loops in intact animals, are sufficient for the generation of coordinated motor patterns (e.g. Bal et al., 1988; Stevenson and Kutsch, 1987). Higher motor centers, on the other hand, have a wide-ranging influence, affecting various body parts concurrently. They may select the appropriate behavior, initiate it, and orchestrate it. To this end, higher motor centers form functional connections with CPGs and may modulate different ones simultaneously. This report describes the interplay between leg CPGs and subesophageal ganglion (SEG), which serves as a higher motor center for locomotion (Gal and Libersat, 2006; Kaiser and Libersat, 2015; Tastekin et al., 2015), of the desert locust, *Schistocerca gregaria* (Forskål).

A comprehensive body of work describing the interactions between higher motor centers and CPGs in the tadpole largely inspired our current work, which shares similar aims with it. Each of the tadpole’s spinal cord segments includes two CPGs that produce an alternating rhythmic motor output, activating the myotomal muscles and thus inducing stereotypic swimming movements (Kahn and Roberts, 1982a). These segmental circuits reciprocally inhibit each other, while maintaining a phase difference between the activity of consecutive segments, thus producing the longitudinal directionality of the movement (Kahn and Roberts, 1982a; b). However, when the spinal cord is isolated from the hindbrain, only short fictive swim bouts can be generated (Li et al., 2006). Further research into these findings revealed that in addition to the spinal network of CPGs, another rhythm-generating center resides in the hindbrain, consisting of electrically coupled descending interneurons (DINs), which are active before each cycle of the spinal motor output, and drive the spinal CPGs (Li et al., 2006, 2009, 2010; Soffe et al., 2009). Similarly, studies in mice and lampreys have pointed to an instrumental role of the brain stem in locomotion (McClellan and Grillner, 1984; Dubuc et al., 2008; Gordon and Whelan, 2008; Hägglund et al., 2010).

To date, the study of higher locomotion centers in insects has mostly focused on their behavioral role *in-vivo*. Using lesions, genetic techniques, and electrophysiological recordings, certain areas of the insect supraesophageal ganglion (brain), such as the central complex and mushroom bodies, were shown to control advanced aspects of walking: for example, speed change and turning (e.g. Strauss, 2002; Gal and Libersat, 2006; Poeck et al., 2008; Bender et al., 2010; Guo and Ritzmann, 2013). The SEG, which anatomically resides between the brain and the thoracic ganglia, was considered responsible for more basic features of walking such as initiation, maintenance, and forward-backward orientation (Huber, 1960; Kien and Altman, 1984; Bässler et al., 1985; Kien, 1990a; Gal and Libersat, 2008; Bidaye et al., 2014). In accordance, lesion-based experiments have demonstrated that removal of the brain does not abolish spontaneous walking, whereas the removal of the SEG eliminates it (Kien and Williams, 1983; Gal and Libersat, 2006). Our knowledge of the anatomical and moreover functional connections between the SEG and leg CPGs is, however, still incomplete.

In accord with previous studies that found central connections between the locust limbs (e.g. Berkowitz and Laurent, 1996), we have recently offered a detailed description of the functional connections among the inter-leg coxa-trochanteral CPGs in the locust *in-vitro* (Knebel et al., 2017). We showed that each of the three thoracic ganglia has its own default, inherent, bilateral coupling of these CPGs: in-phase within the pro- and mesothoracic ganglia, and anti-phase in the metathoracic ganglion. Furthermore, each ganglion was found to be capable of imparting its coupling scheme onto the other ganglia. Importantly, as in most walking systems (Büschges et al., 2011), none of the observed *in-vitro* inter-leg coordination schemes resembled a functional walking gait, and specifically not the tripod gait common among insects, in which neighboring pairs of legs show antiphase activity (Wilson, 1966; Grabowska et al., 2012). The thoracic network of oscillators must, therefore, feature inherent flexibility in order to establish a functional walking gait.

In the current report we follow-up on this work, taking advantage of the locust’s easily accessible nervous systems, as well as its anatomical modularity, i.e. the functional and anatomical separation of the leg CPGs, compartmented by the three interconnected segmental thoracic ganglia; and the higher locomotion centers, encompassed in the two head ganglia – the brain and SEG (or gnathal ganglia; Ito et al., 2014). We introduce first evidence of leverage points for central regulation of the leg CPG network by the SEG higher motor center.

## Materials and Methods

### Experimental animals

All experiments were performed on adult male desert locusts (*Schistocerca gregaria*) from our colony at Tel Aviv University (Ayali and Zilberstein, 2002), within the first two weeks after the final molt. All experiments complied with the Principles of Laboratory Animal Care and the Israeli Law regarding the protection of animals.

### Preparation

Recordings were conducted from *in-vitro* ventral nerve chord preparations, including the pro-, meso- and metathoracic ganglia, and the SEG. Prior to dissection, the animals were anesthetized with CO_2_ for at least 5 min. Following removal of the brain (decerebration), appendages, the pronotal shield, and the abdomen posteriorly to the fourth abdominal segment, a longitudinal cut was performed in the cuticle along the dorsal midline of the thorax. The preparation was pinned to a Sylgard dish (Sylgard 182 silicon Elastomer, Dow Corning Corp.), and the cut was widened and superfused with locust saline containing (in mM): 150 NaCl, 5 KCl, 5 CaCl_2_, 2 MgCI_2_, 10 Hepes, 25 sucrose at pH 7.4. Air sacs and fatty tissue covering the ventral nerve cord were removed, and the thoracic ganglia chain and SEG with their surrounding tracheal supply were dissected out of the animal body, pinned in a clean Sylgard dish, dorsal side up, and bathed in locust saline. The two main tracheae were opened and floated on the saline surface. All peripheral nerve branches originating from the thoracic ganglia were cut short except for the N5a nerves (numbered after Campbell, 1961), which each contain three motor axons: the slow and fast trochanteral depressors and a common inhibitor (Fig. 1; Ds, Df, and CI respectively).

**Figure 1.**
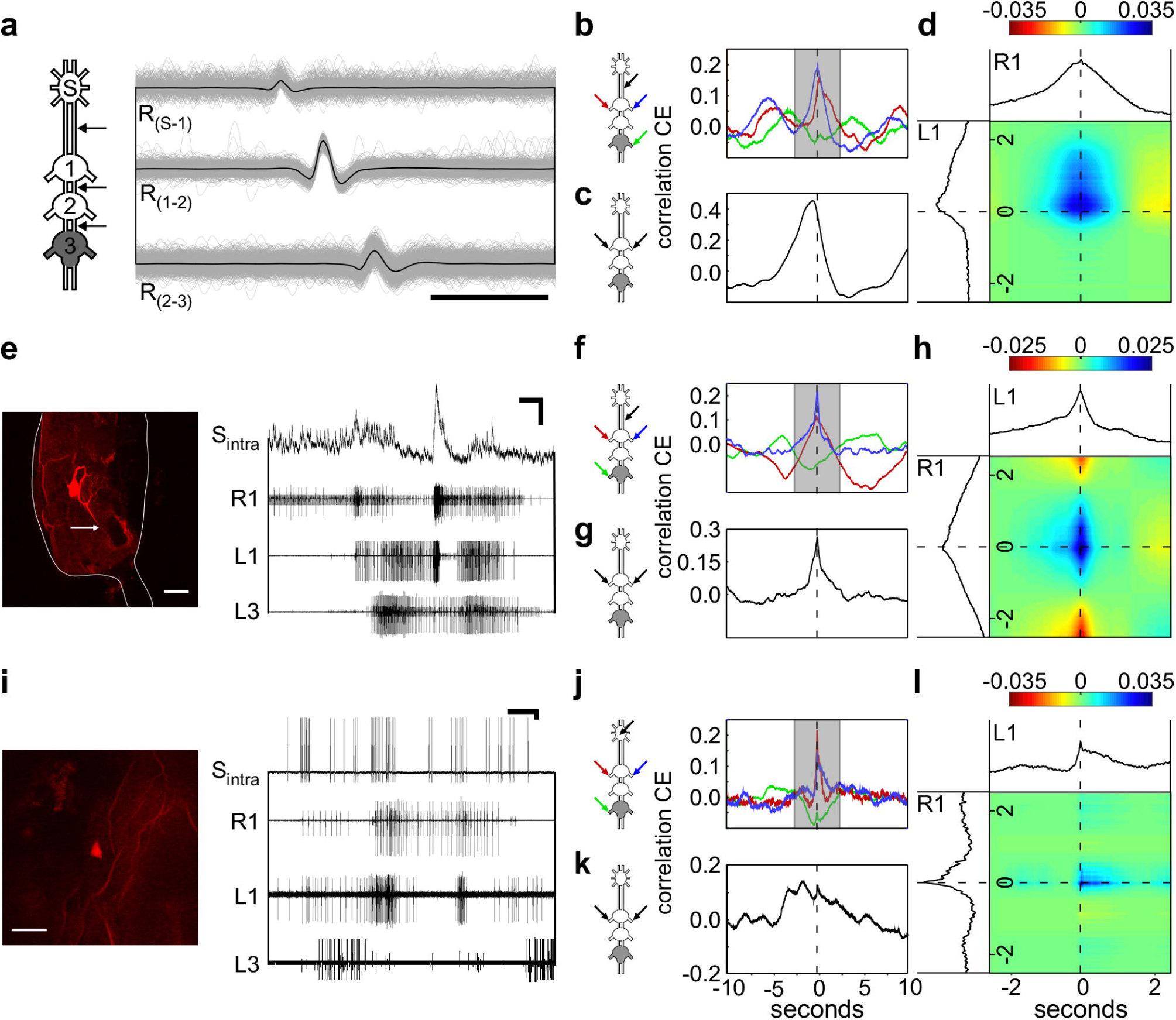
SEG electrical stimulation elicits leg motor outputs. **(A)** Scan of the SEG after staining one neck connective. The dyed somas represent about half of the SEG DIN population. Scale bar: 100 μm **(B)** Maximum intensity projection showing the double labeling of neck connective (red) and nerve 5A staining (green) in the metathoracic ganglion. Arrows in the pictograms and the scan indicate the stained nerves. Scale bar: 100 μm. **(C)** A single horizontal optical section from the framed zone in **(B)**, showing the overlapping area of the depressor motor neurons and DINs projections. Scale bar: 100 μm **(D)** An example of a typical SEG stimulation of the neurons of the mandibular neuromere resulting in a brief elicited activity of the depressor MN of the pro- and mesothoracic ganglia. Scale bar: 5 s **(E)** An example of a typical SEG stimulation of the labial neuromere resulting in a prolonged rhythmic bursting activity of all recorded thoracic ganglia (Df, fast depressor; Ds, slow depressor; CI, common inhibitor). Scale bar: 5 s.

### Electrophysiology and neuroanatomy

To test the effect of SEG electrical stimulation on the thoracic motor output, trains of short electrical pulses (200 ms of 250 Hz pulses of 0.5 ms each, 1 V) were delivered to different areas of the SEG by carefully-fashioned insulated platinum-iridium electrodes, with especially high resistance and small tip diameter (15 μm), as described by Hussaini and Menzel (2013). Thus, we limited the stimulated area as much as possible. All electrical stimulations were generated by a Master 8 stimulator (A.M.P.I).

The activity of the motor nerves (N5a) of each thoracic ganglion was extracellularly recorded by suction electrodes made of borosilicate glass capillaries (A-M systems) and pulled with a P87 puller (Sutter Instruments). Unipolar hook electrodes were used to monitor inter-segmental information transfer from the inter-ganglia connectives (SEG-pro, pro-meso, meso-meta). Data were acquired using two four-channel differential AC amplifiers (Model 1700, A-M Systems).

For intracellular recordings, the SEG neurons were impaled with borosilicate glass capillaries pulled with the P87 puller. The electrode tips were filled with 3 % Neurobiotin (Vector Laboratories, Inc.) diluted in 3M potassium acetate and the shafts with 3M potassium acetate alone, leaving a small air bubble in between. These electrodes had a resistance of approximately 60 MΩ. In order to penetrate the ganglion sheath, a few crystals of protease (Sigma-Aldrich) were placed on the ganglion for 30 s. Intracellularly recorded signals were amplified using a DC amplifier (Axoclamp-2B, Molecular Devices). All recordings were digitized (Axon Digidata 1440A A-D board) and stored on a computer using standard software (Axo-Scope software; Molecular Devices). Resting membrane potentials for all intracellular recordings shown were below −65 mV. After evaluating the responses to restricted drug application, the intracellularly recorded neuron was labeled by iontophoretic injection of Neurobiotin using depolarizing current pulses (2 nA; 100 ms; 5 Hz; 20-40 min).

Backfill staining was accomplished by cutting the desired nerve and bathing it in Dextran-Rhodamine (Molecular probes) or Neurobiotin. After intracellular labeling or backfill staining, preparations were placed for 24 h in a moisture chamber at 4°C to allow the tracer to diffuse along the neurons. Thereafter, the nerve chord was fixed for 2-4 h in 3% paraformaldehyde. The ganglia sheaths were permeabilized with a 0.1% collagenase/dispase (Sigma-Aldrich) solution for 30 min at 36°C before the Neurobiotin labeling was visualized by streptavidin-Cy3 (Jackson ImmunoResearch Labs). Finally, the ganglia chain was dehydrated in an ethanol series and cleared in methyl-salicylate (Merck KGaA). The whole-mount preparations were scanned with a confocal microscope (either ZEISS LSM 510, Carl Zeiss, or LEICA TCS SP2, Leica). Neurons were reconstructed from confocal image stacks using Fiji software (http://fiji.sc/wiki/index.php/Fiji).

### Pharmacological treatment

The muscarinic receptor agonist pilocarpine hydrochloride (Sigma-Aldrich, St Louis, MO, USA) was dissolved in locust saline to a final concentration of 0.5 mM, which typically elicits rhythmic motor activity in leg motor nerves (e.g. Ryckebusch and Laurent, 1993). After 5 min in normal saline, the pilocarpine solution was restrictively bath-applied to the metathoracic ganglion, which was isolated from the rest of the ventral nerve cord by surrounding it with a petroleum jelly (Vaseline) wall. All other thoracic ganglia and the SEG were bathed in normal saline. The well was carefully leak proofed by applying saline, first only to the surrounding of the well, then into the well only, and checking for leaks from the well walls.

### Data analysis

The pilocarpine-induced motor activity in the thoracic ganglia was measured in 31 experiments, initially with the SEG intact and subsequently again after disconnecting the SEG. In each experiment between three to seven simultaneous recordings were conducted from the leg depressor motor neuron (MN) pools, including the slow and fast trochanteral depressors and common inhibitor (Fig. 1; Ds, Df, and CI respectively). Spikes were detected and identified based on their amplitude, and only activity of the excitatory MNs was taken into account, without separating between the Ds and Df. Additionally, in some of the experiments the activity of the inter-ganglia connectives and SEG interneurons was also monitored. We routinely evaluated the activity in two subsequent 8 min windows: the first, typically after 22 min post drug application, and the second immediately after the neck connectives were cut.

To identify SEG DINs in the connective recordings, spikes were first identified using a template recognition function (Dataview software, University of St. Andrews) in one of the channels. Subsequently, we overlaid all identified spikes in 20 ms windows by aligning their maximum point. We repeated this process for the parallel time windows in the other connectives recordings, and averaged each of the overlays. This allowed us to examine the typical activity before and after the identified spike in the more rostral and/or caudal connective recordings. Based on a typical axonal conduction speed of 2 m/s (Gray and Robertson, 1998), and a distance of 3-5 mm between each pair of electrodes, we expected DINs spikes to appear at a delay of about 1.5-2.5 ms between electrodes monitoring adjacent ipsilateral connectives, from rostral to caudal. Spikes that were not accompanied by such preceding or delayed activity in adjacent connectives were filtered out. In one experiment, we used hook electrodes to record both neck connectives and to identify simultaneous bilateral descending spikes. After verifying that the spike was descending (as described above), we selected only simultaneous spikes in both connectives.

To characterize the phases between the output of pairs of CPGs, we used cross-spectrum analysis in MATLAB (MathWorks Inc.), following a procedure developed by Miller and Sigvardt, (1998; see also Sigvardt and Miller, 1998). Only significantly coherent frequencies of each pair of recordings were used to calculate each experiment mean phase vector. The Watson–Williams F-test was used to test for differences in the phase vectors. In order to combine both the uniformity and directionality of the phase distributions, we used the synchronization index, ranging from −1, perfect anti phase, to +1, perfect in phase. This was calculated by averaging all the experiments mean phase vectors, and projecting the product on the 0-180 axis (see Knebel et al., 2017 for more details on the cross-spectrum and synchronization index).

In order to compare the SEG neuronal output and the activity of thoracic MN pools, we utilized cross-covariance analysis in MATLAB. We used a smoothed on-time vector of the spikes of each of the simultaneous recordings. It should be noted that the resultant covariance coefficient value was normalized in order to be able to compare between different analyses. The coefficient is valid only relatively as it is highly dependent on the prior smoothing process: the change of the coefficient over time represents the relative likelihood of spikes to occur in different temporal circumstances, and can be compared among the different examples presented that went through the exact same process.

We tested the statistical significance of the obtained covariance coefficients by performing the exact same analysis on 1000 randomly chosen pairs of the motor output recordings, in which each recording came from a different experiment (different locust). Thereby, we could calculate a bootstrap which represented a confidence interval of 95%, which defined the range of covariance coefficients that could be obtained by chance (<95%). The extreme values of the bootstrap were 1.11 and −0.968. Thus, any result that crosses these have less than a 5% chance of being a type 1 error, and therefore can be considered statistically significant. All the results which are mentioned as meaningful crossed this threshold, unless noted otherwise.

To merge the cross-covariance of the activity of two MN pools and SEG neuronal activity, we multiplied the two normalized cross-covariances calculated separately for each MN pool against the SEG neuronal activity. This yielded a square matrix, in which the axes represent the lags of each cross-covariance. Thus, the diagonal axis (from bottom left to top right) depicts the on-time correlation of the MN pool in respect to the SEG activity, while any deviation from this line indicates the correlation value at a certain time-lag. Again, the absolute values of this analysis are relative, but can be compared among themselves to show the tendency of firing at different temporal states of the network.

## Results

All electrophysiological experiments (N=40) were performed on isolated ventral nerve cords of the locust, including the SEG and the three thoracic ganglia. In all, we evaluated the activity of the coxa-trochanteral CPGs, which induce alternated motor output between the leg levator and depressor motor neuron pools (Ryckebusch and Laurent, 1993; Rillich et al., 2013; Knebel et al., 2017). Based on this robust motor pattern, we measured the activity of trochanteral depressor MNs as representatives of the leg motor output, which participate, among others, in the stance phase during stepping.

### The DINs deliver the SEG descending control of the thoracic motor centers

The SEG higher motor center commands are delivered downstream to the peripheral motor center via a set of DINs (Kien, 1990a; b; Kien et al., 1990; Gal and Libersat, 2006). We performed three experiments in which we stained one neck connective, and found 146, 153, and again 153 stained neuronal somas in the SEG (example: Fig. 1A). Therefore, assuming symmetry, and taking into account the few neurons that send descending neurites in both connectives, there are almost 300 SEG DINs (in accordance with Kien et al., 1990). To explore the anatomical relations between the head ganglia DINs and the leg depressor MNs, we performed a double labeling of the metathoracic N5a nerves and one neck connective. This revealed that many DIN projections overlap with the arborization area of the N5a neurons, both ipsilaterally to the stained connective and contralaterally (Fig, 1B and C). We could not determine whether the DINs originate in the brain or the SEG, or whether they form synapses with the motor neurons. However, the overlapping between DINs’ projections and the leg motor neurons are consistent with the previously reported functional connections, both for the SEG DINs (cockroach: Gal and Libersat, 2008; locust: Kien, 1983; Kien and Altman, 1984) and the brain DINs (cockroach: Bender et al., 2010; Mu and Ritzmann, 2008; Ridgel and Ritzmann, 2005).

### The intact SEG fails to induce activity in the thoracic ganglia chain

Previous studies have shown that headless insects, lacking both the supra- and subesophageal ganglion, do not engage in spontaneous walking, and upon tactile stimulation walk only briefly. However, when the SEG was left intact, the insects tended to walk spontaneously and often uninhibitedly, without any additional stimulation (e.g. stick insect: Bässler, 1983; locust: Kien, 1983; cockroach: Gal and Libersat, 2006). Accordingly, several studies have also reported that without the SEG the leg CPGs are inherently inactive *in-vitro* (Knebel et al., 2017 and references within). Therefore, we first explored whether the intact SEG induces spontaneous fictive leg motor activity in the *in-vitro* thoracic ganglia chain preparations.

In all experiments (N=40) the pro- and mesothoracic ganglia depressor MN pools were silent, whereas the meta-thoracic slow depressor was tonically firing (examples: Fig. 1D and E before the stimulus; see also Knebel et al., 2017; Rillich et al., 2013). No spontaneous motor bursts of action potentials were evident in any of the recordings.

### Stimulation of the SEG labial neuromere is sufficient for generating leg CPGs activity *in-vitro*

Extracellular electrical stimulation of the SEG was previously reported to evoke bouts of walking in semi-intact locusts (Kien, 1990a). Following this report, we sought to explore the effect of electrical stimulation to the SEG *in-vitro*. A pair of fine electrode was positioned into the SEG by way of a micromanipulator, and trains of short electrical pulses were applied, while the thoracic motor output was recorded. Due to the easy accessibility of the isolated preparation, we were able to direct the stimulating electrode to any of the three SEG neuropiles (the mandibular, maxillar, and labial neuromeres, from rostral to caudal). Furthermore, we were able to position the electrode in an area very close to the SEG longitudinal midline, and together with the high resistance small tip electrode used, the possible stimulation of the lateral tracts, in which most axons of brain DINs run directly to the ventral nerve cord, was limited.

Short electrical stimulations were delivered to 2-3 of the SEG neuromeres in four animals. Each trial consisted of 10 stimulations at intervals of 30 sec. In all experiments, a motor response in at least one thoracic ganglion was recorded. However, stimulation of the different neuromeres evoked different motor outputs: The mandibular and maxillar stimulations resulted in short responses (example of a mandibular stimulation: Fig. 1D; overall medians: mandibular 1.4 sec and maxillar 4.1 sec), whereas the labial stimulation elicited prolonged activity with an overall median of 22.9 sec. Moreover, stimulation of the labial neuromere elicited rhythmic bursting activity of up to six bursts, in 2-4 of the recorded thoracic leg CPGs (example: Fig. 1E).

Rhythmic bursting activity in a sensory-deprived preparation is necessarily the product of CPG activation. Our findings thus indicate that the labial neuromere of the SEG is sufficient to activate the leg CPGs in all three thoracic ganglia. Interestingly, during all the *in-vitro* labial neuromere stimulations we observed prothoracic bilateral synchronized excitation, whereas coupling patterns among the other recorded nerves were varied (example: Fig. 1E).

### The SEG has no effect on specific bursting properties of the leg CPGs

The muscarinic agonist pilocarpine is known to activate leg CPGs in isolated nervous systems of arthropods (Chrachri and Clarac, 1987; Ryckebusch and Laurent, 1993, 1994; Ryckebusch et al., 1994; Büschges et al., 1995; Johnston and Levine, 2002; Fuchs et al., 2011, 2012; Rillich et al., 2013; David et al., 2016). Recently, we demonstrated that applying pilocarpine restrictively to each of the thoracic ganglia is sufficient for inducing activity in all leg CPGs, including those of the untreated ganglia (Knebel et al., 2017). To explore a tentative effect of the SEG on the CPG-CPG interactions, we activated the thoracic CPGs by applying pilocarpine to the metathoracic ganglion alone in an *in-vitro* preparation, and compared the motor output of the different depressor MN pools both with the SEG intact and following its removal. Thus, while the generation of rhythmicity arose from the caudal end of the ganglia chain, any SEG modulatory influence had to be sent from the rostral end (pictogram: Fig. 2A).

**Figure 2.**
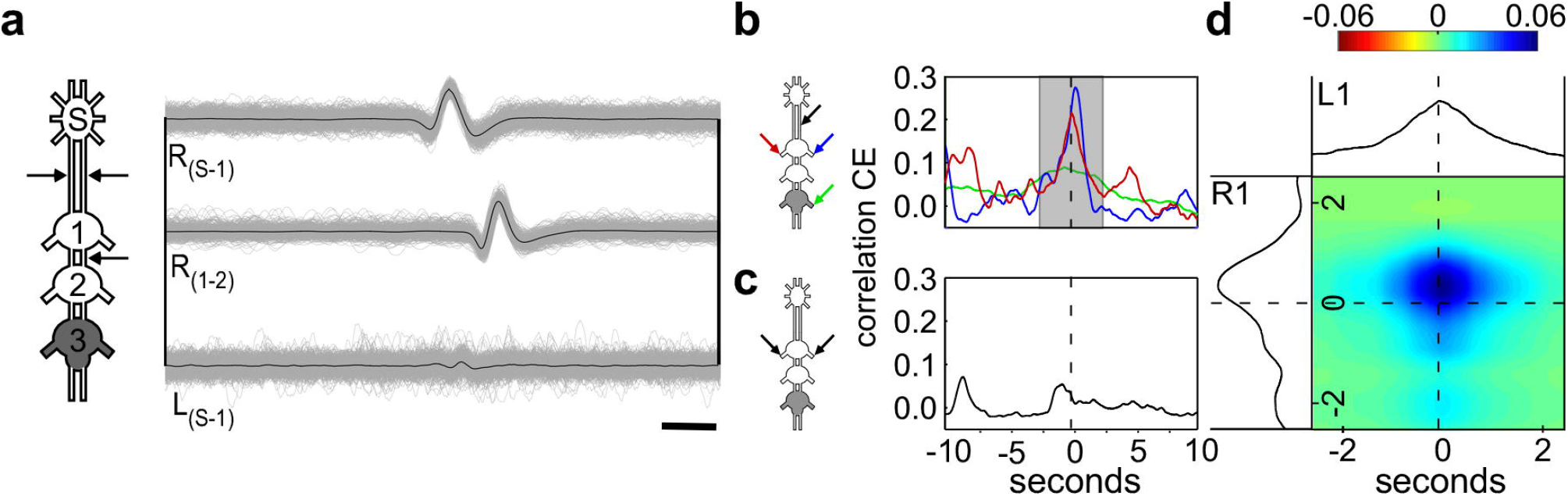
Pharmacological stimulation of the metathoracic ganglion induces rhythmic bursting activity in all three thoracic ganglia. **(A)** The pictogram schematically presents the three thoracic ganglia and SEG. Pilocarpine was applied to the metathoracic ganglion only, as indicated by the gray filling. The arrows indicate the recording sites (4 depressor nerves and 3 connectives), as demonstrated by the recording trace. Scale bar:10 s. **(B)** The spike frequency throughout the 8 min of recording, for each hemiganglion (pro-, meso-, metathoracic hemigagnlia – T1, T2 and T3, respectively), with and without the SEG. No significant difference was found. **(C)** The most common frequency of the pro- and mesothoracic hemiganglia and the ipsilateral source of activation – the ipsilateral hemiganglia (T1 and T2, respectively), before and after SEG removal. Again, no significant difference was found.

Following pilocarpine application, rhythmic motor patterns were obtained in all thoracic ganglia within 2-5 min (Fig. 2A). Before analyzing the coordination pattern among the bursts of the different hemiganglia, we examined whether the SEG influences the general properties of each CPG activity. To this end, we compared the spike frequency in each recording, before and after SEG removal, and found no significant change (Fig. 2B; N=31). We further examined the influence of the SEG on the CPGs recruitment to the pharmacologically activated metathorcic CPGs; as only the metathoracic ganglion was directly exposed to pilocarpine, all other CPG activity resulted from activation of its CPGs. Therefore, the frequencies of the metathoracic CPGs and the other CPGs should resemble each other (i.e. share a common frequency). Using the cross-spectrum analysis, we tested whether the common frequencies differed with and without the SEG (Fig. 2C). We found no effect of the SEG removal on the most common frequencies in the meta-prothoracic and meta-mesothoracic bursting activity (N=31), and therefore concluded that the SEG does not affect the frequency-entrainment of the rostral CPGs by the caudal metathoracic source of rhythmicity. Additionally, we found no SEG effect on the shared frequencies between the two contralateral metathoracic CPGs. Therefore, the SEG has no influence on the rhythmicity of the CPG network.

### The SEG modulates phase parameters among the CPGs

As noted, we have recently reported that in *in-vitro* preparation, when pilocarpine was applied onto the metathoracic ganglion only, all the ganglia bilateral CPGs oscillate in antiphase while the ipsilateral CPGs are synchronized (Knebel et al., 2017). Here we further explored whether the SEG is able to affect this coordination pattern. To this end, we used the synchronization index to determine the synchrony level between each couple of oscillators in our experiments (see Methods for details).

In line with our previous results, when the metathoracic ganglion was directly activated with pilocarpine and the SEG was intact, all ipsilateral CPGs fired bursts of action potentials in-phase. However, unlike in our previous report, in the presence of the SEG the bilateral CPG also oscillated and fired in-phase (Fig. 3C). After removing the SEG, while the ipsilateral CPGs oscillations remained synchronized, the activity of the contralateral CPGs shifted towards anti-phase (Fig. 3D), similar to that recorded in our previous report (where the SEG was removed at the dissection stage; Fig. 3E). Most prominent was the change in the prothoracic ganglion CPGs, whose synchronization index significantly dropped from 0.61 to −0.24 following SEG removal.

**Figure 3.**
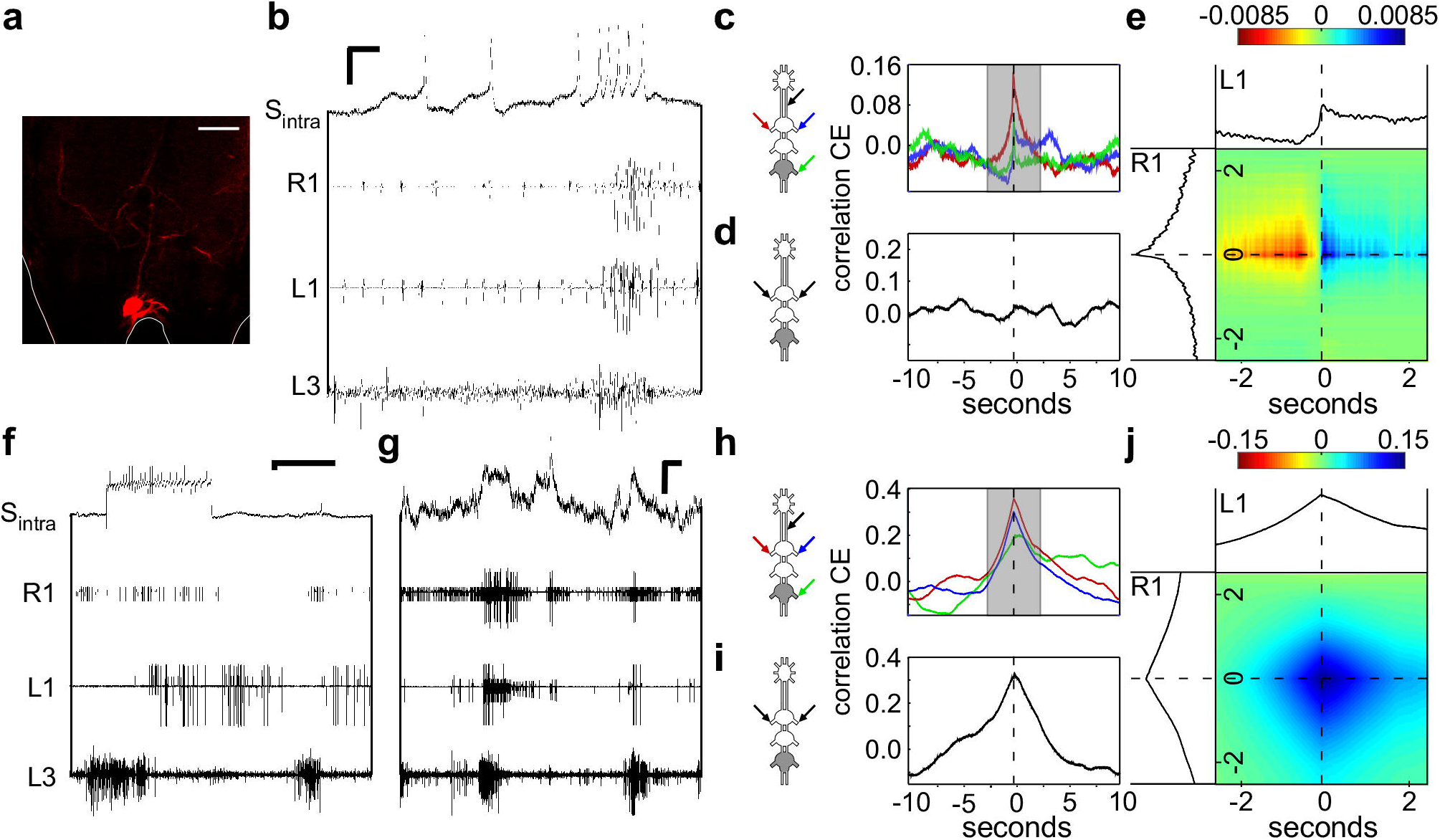
The overall CPG-CPG phase relations are affected by the SEG. **(A)** The bilateral synchrony of the prothoracic CPGs before the SEG removal. The recording exemplifies the typical bursting activity of both hemiganglia. The circular histogram presents the phase of the prothoracic contralateral CPGs of all experiments. The bar filling colors represents the value of the synchronization index, as indicated by the color scale below. The sample size represents only the experiments in which we recorded the bilateral prothoracic activity. **(B)** As in **(A)** but for the activity of prothoracic CPGs after the SEG removal. The phase calculated for the two conditions was significantly different. Scale bar: 2 s. **(C)** A scheme of the CPGs phases before the SEG removal. The colors of the interconnecting lines between the pro-, meso- and metathoracic hemiganglia (upper, middle, and bottom circle pairs, respectively), represent the synchronization index values of each pair of CPGs, as shown on the color scale on the right. **(D)** As in **(C)** but after the SEG removal. **(E)** As in **(C)** but for experiments in which the SEG was removed from the thoracic ganglia chain at the dissection stage. Note that after the SEG removal, the bilateral CPG connections are drawn to anti-phase activity, while the ipsilateral connections remain in-phase, similarly to the experiments in which the SEG was not present from the beginning. *** p<0.001.

Taken together, the SEG synchronizes the activity of the two lateral sides of the CPG network, without affecting the capacity for frequency entrainment and the burst properties of the CPG network.

### Candidate SEG interneurons participate in the bilateral synchronization

To explore the underlying neuronal circuitry behind the bilateral synchronizing effect of the SEG, we simultaneously recorded the CPG motor outputs and the inter-ganglia connectives. Based on multiple connective recordings (example: Fig. 2A) and subsequent spike sorting, we were able to extract the activities of single SEG DINs. Figure 4A-D presents an example of such SEG DIN activity. The cross-covariance analysis shows that this DIN activity was correlated with the firing of the two bilateral MN pools in the prothoracic ganglion: that ipsilateral to the connectives recorded, and that contralateral to them (Fig. 4B). The correlation of the DIN with the latter showed a slight but consistent delay. This difference is visualized in the merged cross-covariance shown in Fig. 4D, where the blue spot, representing a higher correlation between the ipsi- and contralateral prothoracic MN pools in relation to the DIN firings, is smeared towards the positive values of the contralateral prothoracic axis. Since the cross-covariance between the DIN and the metathoracic CPG did not reveal a temporal relationship between their activities (Fig. 4B, green line), this DIN activity is correlated to the activity of specific CPGs.

**Figure 4.**
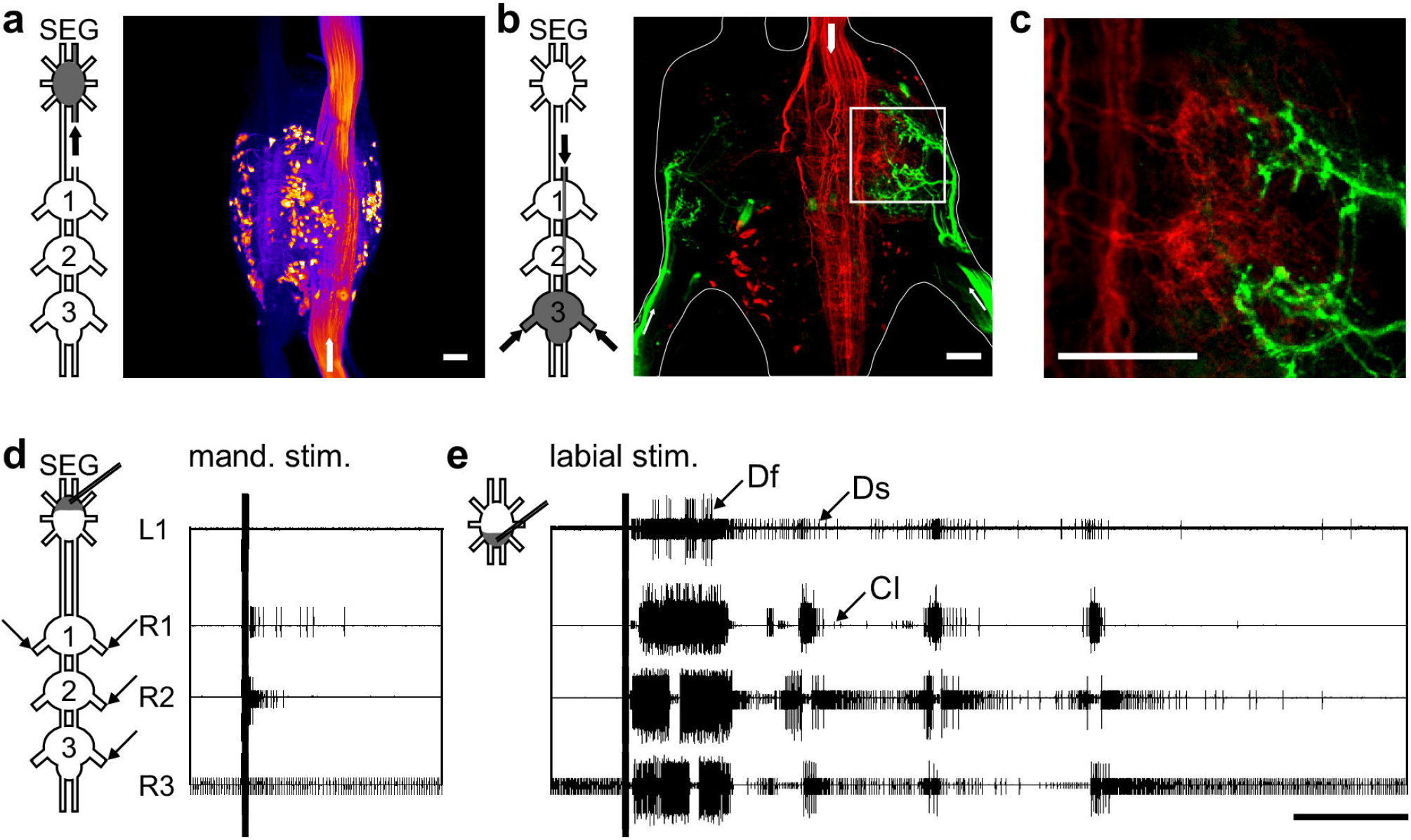
SEG interneurons activity correlates with the prothoracic depressors joint activity. **(A)** Overlays of three simultaneous recordings from the connectives (one for each), indicated by the arrows in the pictogram. The black line represents the average vector of these overlays. The constant delay of 1.8 ms from the SEG downwards to the ipsilateral pro-meso connective indicates that the spike is generated by a neuron in the SEG, sending a descending neurite to all thoracic ganglia. Scale bar: 5 ms **(B)** Cross-covariance between the SEG DIN firing and the left and right prothoracic depressors (blue and red lines, respectively) and the right metathoracic depressor (green line). **(C)** Cross-covariance between the left and right prothoracic CPGs. **(D)** Merged cross-covariance of the left and right prothoracic depressors with the SEG DIN, from the narrow grayed window in **(B)**. The intersection of the 0 values presents the time of the SEG DIN firing, and the colors indicate degree of correlation between the left and right depressors, as indicated by the color scale above. Note that the blue spot in **(D)** is smeared upwards, indicating that the peak of correlation is not at the spike onset, and a delayed prolonged elicited activity is recorded from the CPG contralateral to the DIN. Since no clear correlation is found between the DIN and the metathoracic depressor, as shown in **(B)**, it is a candidate DIN for specifically synchronizing the prothoracic contralateral CPGs. **(E)** Simultaneous intracellular recording of a SEG neuron and the thoracic motor activity. Horizontal and vertical scale bars: 1 s and 10 mV, respectively. The confocal scan shows the recorded SEG neuron. Note that the neuron sends a neurite to the neck connective (indicated by the white arrow). Scale bar:80 μm. **(F)** Cross-covariance between the SEG DIN membrane potential and the left and right prothoracic depressors (blue and red lines, respectively) and the right metathoracic depressor (green line). **(G)** As in **(C)**, but for the prothoracic activity in the preparation shown in **(E)**. **(H)** As in **(D)**, but for the prothoracic and DIN activity in this preparation shown in **(E)** Note the resemblance to the SEG DIN shown in **(D)**. **(I)** Simultaneous intracellular recording of SEG neuron and the thoracic motor activity. Horizontal and vertical scale bars:1 s and 10 mV, respectively. The confocal scan shows the recorded SEG neuron. Scale bar: 80 μm. **(J)** Cross-covariance between the SEG interneuron spikes and the left and right prothoracic depressors (blue and red lines, respectively), and the right metathoracic depressor (green line). **(K)** As in **(C)**, but for the prothoracic activity in the preparation shown in **(I) (L)** As in **(D)**, but for the prothoracic and DIN activity in this preparation shown in **(I)**. Note that despite the weak correlation between the left and right depressors, as seen in **(K)**, the correlation at the time of the SEG neuron spike is clear, as seen in **(J)**.

By means of intracellular recordings we obtained further insights into the activity of SEG interneurons during the metathoracic pilocarpine-induced rhythm. Figure 4E presents an example of a SEG DIN recording. This cell’s activity resembled that of the DIN described above, as can be seen in the cross-covariance analysis: correlated activity with both pro-thoracic MN pools, with a slight difference in the shape of the correlation over time, and only weaker correlation with the metathoracic MNs (Fig. 4F). The recording exhibits rather small action potentials and robust changes in the membrane potential, indicating soma rather than axonal recordings. Hence, to correlate this DIN activity with the prothoracic motor output, we used the values of the membrane potential (and not only the spike on-times).

The two examples presented above demonstrate correlations between the output of both of the prothoracic hemiganglia and an SEG DIN. However, in both cases, the left and right CPGs were rather synchronized even irrespective of the DIN activity (Fig. 4C and G), and it is impossible to distinguish whether the DINs’ activity was connected to one or both the prothoracic CPGs. Therefore, we further analyzed the findings from other experiments, in which the CPG-GPG correlation was lower. Figure 4I presents an example of an intracellular recording of an SEG interneuron in a preparation, in which the prothoracic bilateral synchronization was relatively small (Fig. 4K). Nonetheless, each of the prothoracic CPGs showed correlation with the SEG interneuron (Fig. 4J). Moreover, the merged cross-covariance visualizes the small time window, in which the SEG interneuron was correlated with both prothoracic CPGs (Fig. 4L), in contrast to the overall low bilateral correlation. These findings suggest that this SEG neuron interacts with both prothoracic MN pools.

### SEG descending dorsal unpaired median (DUM) neurons interact with the CPG network

The coupling between independent CPGs can be modulated by neuromodulators such as the biogenic amine octopamine (e.g. Rand et al., 2012; Rillich et al., 2013). To determine whether octopaminergic SEG neurons are centrally coupled to the leg CPGs and are possibly involved in their synchronization, we aimed at specifically recording the octopaminergic SEG DUM neurons.

Due to their unique bilateral descending neurites (Kien et al., 1990; Bräunig, 1991; Cholewa and Pflüger, 2009), we were able to identify the DUM neurons’ activity by recording from both neck connectives, and from the pro-meso connective simultaneously (see Methods for details; Fig. 5A; the different spike amplitudes seen in the left and right connectives in this example is probably a result of different positions of the hook electrode on the two connectives;). Repeating the cross-covariance analysis for this DUM neuron and the two prothoracic CPGs indeed revealed a relatively strong correlation among the activity of all three (Fig. 5B and D). In this example, again, the correlation of each of the prothoracic CPGs with the DUM neuron exceeded the correlation between both CPGs (Fig. 5C).

**Figure 5.**
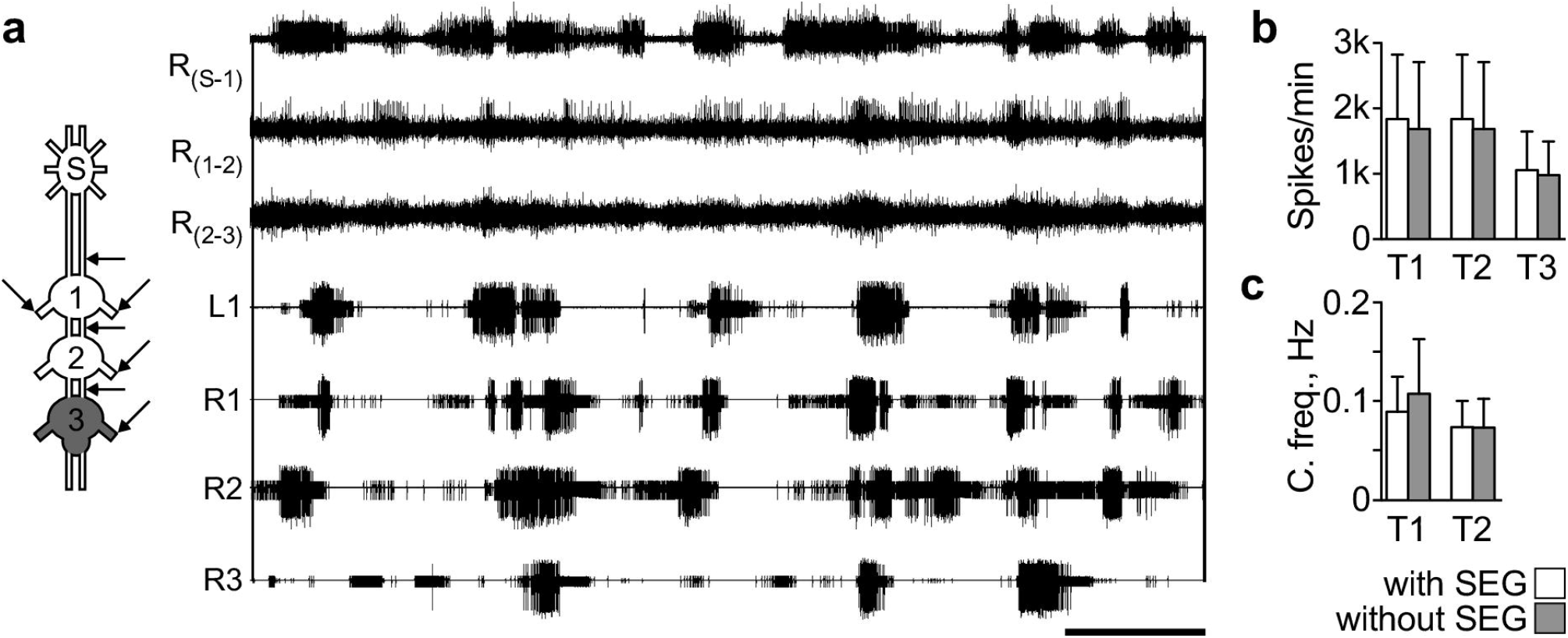
A SEG DUM neuron activity is correlated with the prothoracic depressors joint activity. **(A)** An overlay of 3 recordings from the connectives, shown by the arrows in the pictogram. The almost simultaneous spike (constant delay of 0.4 ms) in both the left and right connectives indicates that the spike is generated by a neuron in the SEG, sending two descending neurites to the thoracic ganglia, typical for SEG DUM neurons. Scale bar: 5 ms. **(B)** Cross-covariance between the SEG DIN firing and the left and right prothoracic depressors (blue and red lines, respectively) and the right metathoracic depressor (green line). **(C)** Cross-covariance between the left and right prothoracic CPGs. **(D)** Merged cross-covariance of the left and right prothoracic depressors with the SEG DIN, from the narrow grayed window in **(B)**. The intersection of the 0 values presents the time of the SEG DIN firing, and the colors indicate degree of correlation between the left and right depressors, as indicated by the color scale above.

Additionally, we intracellularly recorded from a SEG DUM neuron as confirmed by its labeling and action potentials (Fig. 6A): the neuron had a large soma of about 45 μm in diameter, was located medially on the dorsal posterior side of the SEG, and had a symmetrical arborization pattern in the SEG, similar to those described by Bräunig (1991; see laso Kien et al., 1990 and Bräunig and Burrows, 2004). Furthermore, its action potentials were long lasting (~3.5 ms; Fig. 6B), and its soma was excitable (Fig. 6F), as typical for DUM neurons (soma spikes: Heidel and Pflüger, 2006). Similar to the DUM neuron identified by recording the connectives spikes (Fig. 5), this neuron’s action potentials showed a tendency to be synchronized with the output of both prothoracic CPGs (The correlation was statistically significant only with the left nerve recording), while the activity of the CPGs themselves was not correlated (Fig. 6C and D).

**Figure 6.**
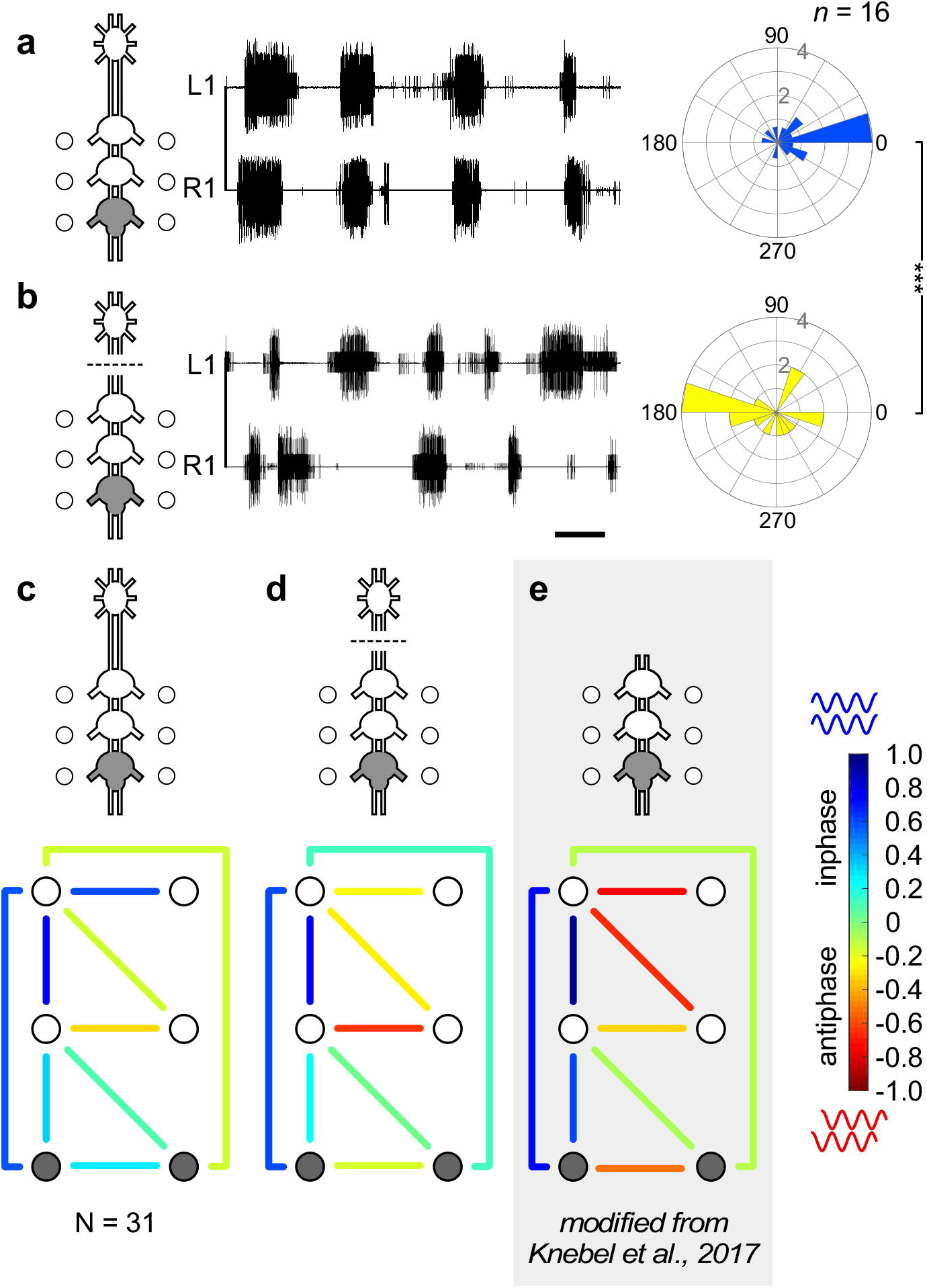
A SEG DUM neuron activity is correlated with the prothoracic depressors joint activity. **(A)** Confocal scan shows the recorded SEG neuron. Scale bar: 80 μm. **(B)** Simultaneous intracellular recording of SEG neuron and the thoracic motor activity. Horizontal and vertical scale bars: 50 ms and 10 mV, respectively. Note the medial position of the neuron, its symmetrical arborization, and its long duration soma spikes. Together, these neuronal characteristics indicate that it is a SEG DUM neuron. **(C)** Cross-covariance between the SEG DIN firing and different sets of depressors (right prothoracic: blue; left prothoracic: red; right metathoracic: green). **(D)** Cross-covariance between the left and right prothoracic depressors **(E)** Merged cross-covariance of the left and right prothoracic depressors with the SEG DIN, from the narrow grayed window in **(C)**. The intersection of the 0 values presents the time of the SEG DIN firing, and the colors indicate degree of correlation between the left and right depressors, as indicated by the color scale above. **(F)** Simultaneous intracellular recording of SEG neuron and the thoracic motor activity before, during, and after a short depolarization. Horizontal and vertical scale bars: 2 s and 10 mV, respectively. **(G)** Simultaneous intracellular recording of the SEG DIN and the thoracic motor activity during a permanent hyperpolarization. Horizontal and vertical scale bars: 2 s and 10 mV, respectively. **(H), (I)** and **(J)** are the same as **(C)**, **(D)**, & **(E)** respectively, but for the SEG DIN during the permanent hyperpolarization, and conducted by analyzing the changes in its membrane potential.

In order to uncover the postsynaptic potentials (PSPs) that this neuron receives, we hyperpolarized it for several minutes and thus minimized the action potentials generated. We found that the DUM neuron received accurate and consistent information about co-activation of the two prothoracic CPGs (example: Fig. 6G; analysis: Fig. 6H). Since the DUM neuron EPSPs indicate synchrony between the bilateral CPGs, they might offer a mean of coincidence detection. Interestingly, during the period of hyperpolarization, the synchronization among the prothoracic CPGs noticeably increased (Fig. 6I). Depolarization, on the other hand, did not evoke any immediate response (Fig. 6F).

## Discussion

In this study we have demonstrated two major aspects of functional interactions between the higher locomotion centers of the SEG and the leg CPGs in the locust: (1) the ability of the SEG to induce CPG activity, and (2) the role of the SEG in coordinating the coupling among the leg CPGs. These interactions are mediated by the SEG DINs, connecting the SEG with the thorax.

### The use of *in-vitro* preparations

All the experiments presented here were conducted *in-vitro*. As previously explained in Knebel et al. (2017), and as a common working hypothesis in CPG research (e.g. Ayali and Lange, 2010 for review), it is advantageous to study the central interplay of CPGs in the complete absence of sensory inputs. This reductionist approach is consistent with the very definition of CPGs as a neuronal oscillator capable of performing its tasks with no sensory regulation (Marder and Bucher, 2001). However, there are also some clear limitations to this approach: namely, that conclusions, valid at the level of the neuronal network and nervous system, may not be directly reflected in the actual behavior, which is further shaped by additional sources, such as sensory inputs and the animal’s internal state, in the intact animal. Nonetheless, it is important to note that since the central connections and interactions we studied here *in-vitro* are those underlying the execution of the motor behavior, the insights gained *in-vitro* are instrumental for the generation of hypotheses and predictions regarding *in-vivo* insect locomotion.

### The SEG’s ability to initiate different activity patterns in leg CPGs

By means of electrical stimulation of SEG neurons we were able to activate each of the leg coxa-trochanteral CPGs. This finding, obtained in the isolated *in-vitro* preparation, indicates the sufficiency of the central connections between the SEG and the thoracic ganglia in activating the leg CPGs. Moreover, of the three SEG neuromeres, overall containing approximately 300 SEG DINs (Fig. 1A; see also Kien et al., 1990), we identified the labial neuromere as the most potent SEG area for CPG activation. Labial neuromere excitation induced prolonged bursting rhythms, similar to those obtained by activating the leg CPGs pharmacologically (compare examples in Fig. 1E and Fig. 2A; see also Knebel et al., 2017). These results are in accord with those reported by Kien (1983), as well as with our own preliminary experiments (not described herein), showing that stimulating the SEG in a semi-intact preparation results in leg movements (suppl. videos 1&2). Altogether, these findings suggest that the SEG is capable of inducing leg CPGs activity, and thus initiating stepping behavior. Moreover, they also suggest that by providing rhythmical excitatory and inhibitory signals to the leg CPGs, the SEG can maintain ongoing legged activity, as was previously suggested by Kien and Altman (1992). However, we also observed that intact connections to the unstimulated SEG were not sufficient by themselves to initiate spontaneous leg motor activity. It therefore seems that rather than constituting an autonomous “motivation center”, the SEG functions as a relay station or a mediator, and depends on central inputs from the brain, from other CPGs, or on sensory information, in order to drive leg CPGs, (see also Kien and Altman, 1984). Moreover, the motor output induced by the SEG *in-vitro*, was monitored in all the leg CPGs but did not resemble any functional walking gait. This indicates that there is no complete “coordination program” or “walking motor program” conveyed by the SEG to the CPG network. Nonetheless, throughout our experiments, a stereotypic bilateral synchronization pattern of the leg CPGs was induced, suggesting that the SEG can modulate the overall coordination of the locomotion network.

We have tried to limit the likelihood of stimulating severed axons, which originally traveled from the brain to the SEG or thoracic ganglia, as explained in the methods and results sections. However, as with any extracellular stimulation, we cannot rule out the possibility that undesired neurites were stimulated. Yet, our finding that the stimulation of different neuromeres resulted in differentiated activation of the legs MN pools, and that the coordination pattern established resembled that induced by the unstimulated SEG, indicates that at least to the degree of affecting the leg motor output, stimulation of brain DINs did not occur.

### The SEG bilateral synchronizing influence

We characterized the role of the SEG in modulating the output of the leg CPGs network using an *in-vitro* preparation that included the thoracic ganglia and the SEG. Based on our previous study (Knebel et al., 2017), we chose a specific paradigm of restrictive pilocarpine application to the metathoracic ganglion. This allowed us to investigate the output of the more rostral pro- and mesothoracic ganglia, their interconnections, and their connection with the SEG, in the absence of any direct effects of pilocarpine.

We found that the SEG does not alter the firing properties of any individual CPG, nor their mutual bursting frequencies (Fig. 2B and C, respectively) but, rather, selectively affects the phase relations of the bilateral CPG couples, most consistently those of the prothoracic ganglion (Fig. 3). It is possible that the SEG can strengthen the inherent synchronized bilateral coupling of the prothoracic CPGs (Knebel et al., 2017), and the prothoracic ganglion, in turn, induces this coupling pattern upon its neighbors through the synchronizing ipsilateral connections. However, from the neuronal network perspective, since the ability of all the ganglia to induce activity in the CPGs of other ganglia and influence their phase is equal (Knebel et al., 2017), it is more plausible that the SEG modulates all bilateral inter-CPG connections, and that of the prothoracic ganglion is simply the most responsive. Moreover, the SEG had no noticeable effect on coupling among the ipsilateral CPGs, whose activities are synchronized independently of it. It should be noted that these findings were obtained without the animal’s brain, and future studies will investigate its role in the CPGs coordination.

Supporting the findings described above, we found various examples of SEG cells (Fig. 4–6) whose activities increased in phase with the synchronized output of the left and right prothoracic CPGs. In some of these examples, the correlations of the left and right depressors with the SEG neurons were stronger than their mutual correlation (Fig. 4I–L, 5–6), suggesting that SEG neurons play an active role in bilateral synchronization. However, due to the unexcitable properties of the soma membrane of those interneurons, we were unable to elicit action potentials in most of our intracellular experiments, and thus could not reach any causative conclusions regarding their role in the network.

It should be noted that due to our experimental procedure of restrictive pilocarpine application, all correlated activity of SEG interneurons with the leg CPGs must be the result of ascending information reaching the SEG and mirroring the CPG activity. Such information can be referred to as “efference copy” (Jeannerod and Arbib, 2003), and is independent of sensory information (which was unavailable in our experiments).

### The octopaminergic system interplay with the coordination of leg CPGs

A potential candidate for the modulation of CPG-CPG interplay in insects is the octopaminergic system (e.g. Rand et al., 2012; Rillich et al., 2013) and its prominent members, the DUM neurons (for review: Libersat and Pflueger, 2004). Indeed, we found that the activity of a descending SEG DUM neurons was correlated with the bilaterally synchronized activity of the contralateral prothoracic CPGs (Fig. 5–6). This suggests involvement of the octopaminergic system in the inter-CPG coordination, as all DUM neurons in the SEG are octopaminergic (Bräunig, 1991; Stevenson and Sporhase-Eichmann, 1995). Furthermore, we found that EPSPs recorded from a DUM neuron reliably reflected the co-activation of the front leg depressors. Therefore, an ascending neuron, or neurons, form synapses on the DUM neuron, eliciting changes in its membrane potential correlated with the thoracic CPGs activity. This further confirms our claim that a central efference copy of leg activity is delivered to the SEG.

Surprisingly, we found that upon silencing this DUM neuron, the prothoracic bilateral synchronization increased. Taken together with the rest of our findings, this result suggests that different neuronal elements in the SEG coordinate leg activity in a complex, and not necessarily consistent, manner.

### The synergy of leg CPGs, higher motor centers, and sensory information in walking

Walking behavior is dependent on the synergies of CPGs activities, sensory inputs, and higher motor center regulation. Our results indicate the SEG as a potential integration center: receiving descending inputs from the brain, as the anatomy of its neurons and those of the brain suggests (Roth et al., 1994), and ascending information from both leg sensory organs (Kien and Altman, 1984) and the CPGs themselves, as presented here.

Since we did not observe in any of our *in-vitro* experiments fictive walking, namely, a motor output that resembled a functional walking gait, we have to conclude that, in the locust, functional inter-leg coordination during walking is only accomplished by way of complementary sensory inputs or descending commands from the brain. This was also reported for many other walking systems (Büschges et al., 2011) and vast evidence suggests that sensory inputs plays a major role in walking coordination (e.g. Borgmann et al., 2007, 2009; Daun-Gruhn, 2011; Fuchs et al., 2012).

We did, however, find that the SEG affects the overall leg CPG coordination. Büschges et al. (1995; based on v. Holst, 1936) suggested that synchronized activity of locomotive oscillators is the energetically cheapest way to couple them while ensuring their joint frequency. As we have previously shown, the bilateral coupling among the CPGs in the thoracic ganglia is flexible and rather weak compared to the ipsilateral coupling (Knebel et al., 2017). It is therefore plausible that the level of bilateral synchronization mediated by the SEG is sufficient to drive the two ipsilateral trios of CPGs to oscillate at similar frequencies, as required for walking straight for example. Previous studies have already suggested that the SEG is a bilateral mediator (Kien and Altman, 1984; Kien et al., 1990; Bräunig and Burrows, 2004), but this was posited mostly due to the anatomy of the majority of the DIN branches, which are contralateral to their soma (Gal and Libersat, 2006). In light of this known DINs anatomy, and our physiological findings, it is plausible that a decussation of information and commands occurs in the SEG: ascending information about the CPGs activity and leg sensory inputs is received ipsilaterally, while the DINs deliver commands downstream via the contralateral descending axons, thus regulating the bilateral CPGs connections.

### A comparison to vertebrate systems

A comparison of the anatomy and function of the SEG and the vertebrate brainstem suggests some parallels, as previously suggested by Schoofs et al. (2014). In the hindbrain-lesioned tadpole, for example, only short fictive swimming episodes could be elicited in response to tactile stimulus (Li et al., 2006), similar to the short sequences of walking that SEG-less insects perform (Gal and Libersat, 2006). Furthermore, subpopulations of brainstem neurons mapped functionally and anatomically were shown to be involved in locomotion initiation and halt (Shik et al., 1969; McClellan and Grillner, 1984; Sirota et al., 2000; Cabelguen et al., 2003; Hägglund et al., 2010; Esposito et al., 2014; Bouvier et al., 2015). Specifically, in all the different species examined, stimulation of the mesencephalic locomotor region (MLR) induced locomotion-related activity both *in-vivo* and *in-vitro* (Esposito and Arber, 2016). In the tadpole, neuronal pacemakers in the hindbrain that send descending neurites to the swimming CPGs fire at the beginning of each swimming cycle, presumably thereby maintaining a swimming pattern (Soffe et al., 2009). Our experiments showed that the SEG can elicit several burst cycles in the different leg depressors. It is therefore possible that, in a similar manner to the tadpole hindbrain, repetitive activity of SEG DINs at the appropriate time during the step cycle can excite the CPGs to be entrained in a functional manner.

Certain findings have also suggested that the brainstem mediates activity of both sides of the vertebrate body, analogically to the role of the SEG reported here. Each of the two-sided MLRs is capable of eliciting bilateral activation of the locomotion circuits along the spinal cord by activating both sides of the reticulospinal tracts simultaneously, thus delivering symmetrical commands to the spinal cord (Brocard et al., 2010). Furthermore, it is within the medulla oblongata that the pyramidal tracts (which include the motor corticospinal fibers), intersect to deliver information to the contralateral lower motor neurons. Moreover, similar to the brainstem, out of which the cranial nerves emerge, the SEG control the insect’s mouthparts.

## Conclusion

The SEG plays a regulative role in control of the arthropod legs. It has the ability to induce movement in all legs in different coordination patterns by activating their CPGs, and it mediates their bilateral activity by adjusting the CPG coupling. Our findings have revealed ways by which the SEG higher motor center serves as a higher motor center of legged activity, as has been suggested in previous studies (Kien, 1983; Gal and Libersat, 2006; Bidaye et al., 2014). Overall, we show that the SEG interacts with the leg CPGs to organize their joint activity; and, being situated in-between the brain and thorax, it bridges between the brain centers, sensory inputs, and the motor circuits of the legs. Interestingly, a comparison between the SEG and the vertebrate brain stem suggests some parallels.

As noted above, findings based on *in-vitro* preparations are sometimes difficult to interpret, especially when they do not completely correspond to the real behavior. However, our findings did reveal functional connections between the leg CPGs and the SEG. Hence, the importance of the results, for understanding walking behavior, is clear. In light of the current findings and those of our previous study (Knebel et al., 2017), it is now known that the locust CPGs, with and without the SEG inputs, are not naturally coupled to produce any walking-like coordination. This is a consequence of the highly modulated system that has to adjust to the heterogeneous environment in which walking occurs, and therefore must avoid predetermined deterministic movements. Future work should explore the effects and the roles of other mechanisms modulating the CPG network, such as further higher motor centers and sensory inputs, and assess their contribution to walking.

## Acknowledgements

This work was partially supported by DAAD travel scholarships (to DK) and, in its final stages, by the German Research Council (DFG; Grant RI 2728/2-1). H-J P gratefully acknowledges the support by the DFG (FOR 1363, Pf128/30-1 and PF128/32-1) and the receipt of a Berlin NaFoeG-stipend to LN. We thank Baruch Barzel for his valuable support in the data analysis.

